# The cooperative assembly of shelterin bridge provides a kinetic gateway that controls telomere length homeostasis

**DOI:** 10.1101/2020.07.31.231530

**Authors:** Jinqiang Liu, Xichan Hu, Kehan Bao, Jin-Kwang Kim, Songtao Jia, Feng Qiao

## Abstract

Shelterin is a six-proteins complex that coats chromosome ends to ensure their proper protection and maintenance. Similar to the human shelterin, fission yeast shelterin is composed of telomeric double- and single-stranded DNA-binding proteins, Taz1 and Pot1, respectively, bridged by Rap1, Poz1, and Tpz1. The assembly of the proteinaceous Tpz1-Poz1-Rap1 complex occurs cooperatively and disruption of this shelterin bridge leads to unregulated telomere elongation. However, how this biophysical property of bridge assembly is integrated into shelterin function is not known. Here, utilizing synthetic bridges with a range of binding properties, we find that synthetic shelterin bridge lacking cooperativity requires a linker pair that matches the native bridge in complex lifespan but has dramatically higher affinity. We find that cooperative assembly confers kinetic properties on the shelterin bridge allowing disassembly to function as a molecular timer, regulating the duration of the telomere open state, and consequently telomere lengthening to achieve a defined species-specific length range.

## INTRODUCTION

In most eukaryotes, telomeres, the natural ends of chromosomes, are essential for stable maintenance of chromosomes, and thus our genetic information (de Lange, 2018). Similar to humans, the telomere structure of fission yeast, *Schizosaccharomyces pombe*, is achieved by association of shelterin components with both double-stranded (ds) and single-stranded (ss) telomeric DNA, forming a nucleoprotein complex (Miyoshi et al., 2008). Fission yeast Taz1 (TRF1/2 in humans) and Pot1 (POT1 in humans) specifically bind to telomeric double-stranded (ds) and single-stranded (ss) DNA, respectively (Baumann and Cech, 2001; Cooper et al., 1997). In fission yeast, Rap1, Poz1, and Tpz1 bridge the telomeric dsDNA binder Taz1 and ssDNA binder Pot1 through their direct protein-protein interactions forming the Taz1-Rap1-Poz1-Tpz1-Pot1 complex (Fig. 1A). Telomeres are maintained at a species-specific length range and this telomere length homeostasis is proposed to be regulated via dynamic switching of telomeres between two states: telomerase-extendible (open) and telomerase-nonextendible (closed) states. Genetic deletions of shelterin components such as Taz1(Cooper et al., 1997), Rap1 (Kanoh and Ishikawa, 2001), Poz1 (Miyoshi et al., 2008), or mutations that disrupt the connectivity in the shelterin bridges (Rap1-Poz1-Tpz1) lead to drastically elongated telomeres due to unregulated telomerase action on telomeres (Jun et al., 2013).

**Figure 1.**
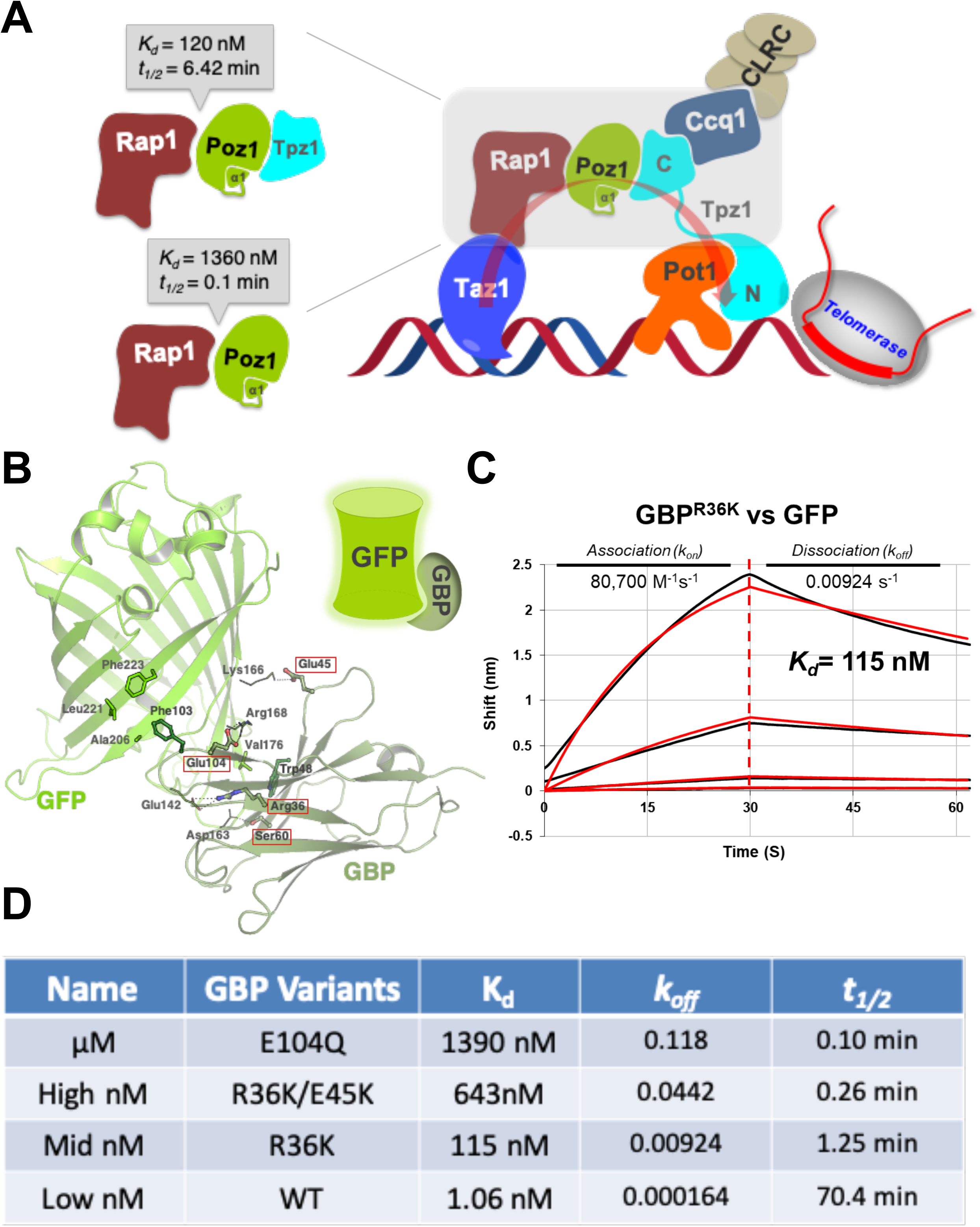
Construction of synthetic shelterin bridges utilizing GFP-GBP pairs of variable binding properties. (**A**) Schematic diagram of *S. pombe* shelterin complex (right) and Tpz1-Poz1-Rap1 interaction (left). <u>*Right*</u>: Overview of *S. pombe* shelterin complex. Rap1, Poz1, and Tpz1 connect double-stranded and single-stranded telomeric DNA binding proteins, Taz1 and Pot1, respectively, forming the shelterin bridge through their protein interactions. Telomerase and histone H3K9 methyltransferase Clr4 complex (CLRC) are recruit to telomeric DNA through shelterin components. For clarity, only one copy of each component is shown, which does not represent the stoichiometry of shelterin complex in cells. <u>*Left*</u>: Binding affinity and half-life calculated from dissociation constant (*k*_off_) of free or Tpz-bound form of Poz1 to Rap1 as reported previously. (B) Structure of GFP-GBP complex and the interface residues. Residues selected in this study to mutate are highlighted with red boxes. (**C**) Bio-layer interferometry (BLI) measurement of dissociation and association events in real time between GFP and GBP variants using Octet red96. A 1:1 binding model is used to fit binding curves globally, yielding equilibrium dissociation constant (K_d_), association (*k*_on_), and dissociation (*k*_off_) rate constants. BLI experiments were repeated three times and representative results were shown. (D) Summary of binding properties for GFP and GBP variants interactions. Half time (t_1/2_) of the protein interaction is calculated using t_1/2_ = ln2/*k*_*off*_.

Our previous work showed that the Tpz1-mediated complete linkage within the shelterin bridge, rather than individual components *per se*, defines the telomerase-nonextendible state (Jun et al., 2013). Moreover, Tpz1 physically interacts with both positive and negative regulators of telomere length and executes its functional roles to coordinate shelterin and telomerase in response to cell cycle signals (Hu et al., 2016; Liu et al., 2015). Through its interaction with Ccq1, Tpz1 also recruits the Clr4 methyltransferase complex CLRC to telomeres and establishes subtelomeric heterochromatin (Wang et al., 2016). Therefore, shelterin bridge has been the emerging key player in regulating telomere length homeostasis, telomeric silencing, and possibly other telomeric functions.

Shelterin complex, like other macromolecular complexes, such as ribosomes, proteasomes, needs to assemble and turn into their functional forms in a timely and precise manner in response to cellular signals. Cooperativity is a general strategy that allows multiple components to rapidly and accurately form a higher-order complex with functional conformation (Williamson, 2008). The fission yeast shelterin bridge, Tpz1-Poz1-Rap1 complex, has recently been shown to assemble cooperatively (Kim et al., 2017). The assembly pathway of this three-component complex was revealed at the atomic resolution. Tpz1-Poz1-Rap1 complex assembly is initiated by the binding of Tpz1 to Poz1, which induces the folding of the N-terminal helix in Poz1, called “conformational trigger”. Consequently, formation of new hydrophobic interactions and hydrogen bonds within Poz1 ensues, which result in conformational changes at the Rap1-binding surface of Poz1, enhancing the binding affinity (K_d_) of Poz1-Rap1 interaction 10 times and decreasing its disassociation rate (*k*_*off*_) 60 times (Fig. 1A). Loss of the conformational trigger causes breakdown of shelterin bridges on telomeres, and leads to unregulated telomere elongation, indicating the essential role that cooperative assembly plays in telomere function. However, how this biochemical feature is integrated into shelterin function or regulation is not known.

In this study, utilizing the well-studied GFP and GFP nanobody (also called GFP Binding Protein or **GBP**) pair (Kubala et al., 2010), we engineered a spectrum of GBP variants that interact with GFP with different thermodynamic and kinetic properties. We then physically tethered the GFP-GBP variant pairs to shelterin components and tested their ability to rescue shelterin bridge defective in cooperative assembly. To rescue telomere length, synthetic shelterin bridge without cooperativity requires a GFP-GBP pair with 30-fold higher binding affinity (K_d_) than that of the native shelterin bridge. Interestingly, this synthetic shelterin bridge recapitulates the assembly kinetics of the native shelterin bridge, having similar dissociation rate (*k*_*off*_) and thus half-life (*t*_*1/2*_). Indeed, our previous work indicates that the assembly of shelterin bridge (Tpz1-Poz1-Rap1 complex) is promoted by the Tpz1-Poz1 interaction, which enhances Poz1-Rap1 interaction mainly by decreasing the disassociation rate of Rap1 from Tpz1-bound Poz1. Therefore, cooperative assembly installs a “kinetic gateway” in the shelterin bridge that controls timespan of the formation-and-breakage of the shelterin bridge, which in turn regulates telomere length. In contrast, telomeric silencing function of shelterin bridge is less dependent on the kinetics of its assembly, but more on its binding affinity, agreeing with its role in passively recruiting and enriching the histone H3K9 methyltransferase Clr4 to telomeres (Wang et al., 2016).

## RESULTS

### Design synthetic shelterin bridge with GFP-GBP pairs of variable binding properties

Recent genetic, biochemical and structural studies utilizing model organism fission yeast, *Schizosaccharomyces pombe*, have uncovered that the shelterin bridge connecting telomeric dsDNA and ssDNA controls the extendable and non-extendable states of the telomeres (Jun et al., 2013). In addition, the integrity of shelterin bridge is also required for telomeric heterochromatin formation (Wang et al., 2016). Crystal structures of fission yeast shelterin bridge (Tpz1-Poz1-Rap1 complex) has provided atomic views of the shelterin bridge and revealed cooperative assembly as the fundamental principle of shelterin bridge formation (Kim et al., 2017). In this process, Tpz1-Poz1 interaction induces conformational changes in Poz1, which greatly enhances the binding of Rap1 to Poz1, forming a stable shelterin bridge. To elucidate the biological significance of the cooperativity in the shelterin bridge assembly, we aimed to interrogate whether the allosterically enhanced thermodynamic binding affinity (K_d_) or the kinetic stability (*k*_off_) of the shelterin bridge originated from Tpz1-induced conformational changes of Poz1 is the key factor in regulating proper telomere length control and telomeric silencing.

To deconvolute the purpose of employing cooperative assembly mechanism for shelterin bridge, we constructed synthetic shelterin bridges utilizing protein-protein interaction module GFP-GBP with variable binding properties. The GFP Binding Protein (GBP) is a GFP nanobody, a single-chain V_H_H antibody domain developed to bind to GFP with high specificity (Kubala et al., 2010). The wide-type GBP interacts with GFP with a thermodynamic dissociation constant (K_d_) of 1.06 nM, which is over 113 times stronger than the binding of Tpz1-bound Poz1 to Rap1. Moreover, the GFP-GBP complex also has very slow disassociation rate with *k*_off_ = 1.64 × 10^−4^ s^−1^, thus the half-life (the time in which half of the initially present complexes have dissociated—1n2/*k*_off_) being 70.4 min, about 10 times longer than that of the Tpz1-bound Poz1-Rap1 complex (6.42 min) and 700 times longer than that of the free Poz1-Rap1 (0.1 min) (Fig. 1B). To construct synthetic shelterin bridges with a spectrum of thermodynamic and kinetic properties, we set out to design a series of GBP mutants that have decreased interaction affinity with GFP based on the crystal structure of GFP-GBP heterodimer complex. GBP-GFP interaction is mostly driven by hydrophobic interactions between F102 and L221, A206 of GBP and F223 of GFP, as well as between W47 of GBP and V176 of GFP. Around the hydrophobic core, GBP forms salt bridges with GFP via its R35, E45 and E103 to enhance specificity and affinity (Fig. S1). To weaken, rather than disrupt GFP-GBP interaction, we selected residues around the edge, but not in the hydrophobic core of the GFP-GBP interface (highlighted residues in Fig. S1). Thus, we introduced mild changes to GBP by mutating these residues to amino acids with similar properties (Fig. 1C). We aimed to select for GBP mutants that have a range of binding affinities with GFP (low nM, middle nM, high nM, and μM) and carried out binding assays using Bio-layer interferometry (BLI) to measure both the thermodynamic binding affinity (K_d_) and kinetic behavior (*k*_off_) of the GFP-GBP variant pairs. As shown in Fig. 1C, 1D, and S1B we successfully achieved this goal by obtaining GFP interacting GBP variants— GBP^R36K^ (115 nM: mid nM), GBP^R36K/E45K^ (643 nM: high nM) and GBP^E104Q^ (1.39 μM: μM). Together with GBP^WT^ (1.06 nM: low nM), we have GFP-GBP pairs with their binding affinities (K_d_) ranging from 115 nM to 1.39 μM levels, and disassociation constant (*k*_off_) ranging from 0.00016 s^−1^ to 0.12 s^−1^ (correspondingly the converted half-life *t*_*1/2*_ from 70.4 min to 0.1 min). Among them, GBP^R36K^ binds to GFP with a similar affinity as Tpz1-bound Poz1-Rap1, but shorter half-life (1.25 min vs. 6.42 min). GBP^E104Q^ binds to GFP with a similar affinity and half-life to Poz1-Rap1 interaction. GBP^R36K/E45K^ lies between the above two GBP variants. GBP^WT^ can provide extremely high affinity and long half-life (70.4 min). With these GBP variant-GFP pairs, we were able to engineer shelterin bridge with synthetic bridges of a wide range of binding properties to investigate which biochemical properties of shelterin bridge enabled by cooperative assembly are determinants of telomere length regulation.

### Slower disassociation rate rather than increased binding affinity is the key contribution of cooperative shelterin bridge assembly to telomere length regulation

The cooperative interaction between Rap1 and Poz1 induced by Poz1-Tpz1 interaction has been elucidated at atomic level in our previous study (Kim et al., 2017). Poz1-R218E was identified as a mutant defective in Poz1-Rap1 interaction due to the disruption of a major salt bridge between Poz1-R218 and Rap1-E476. Moreover, deletion of the extended Poz1-interaction domain of Rap1 (Rap1ΔPID) also results in defective Poz1-Rap1 interaction. Importantly, both Poz1-R218E and Rap1ΔPID mutants selectively disrupt the Poz1-Rap1 interaction without affecting other interactions within the shelterin bridge. Therefore, in either Poz1-R218E or Rap1ΔPID background, physically linking Poz1 and Rap1 using identified GBP-GFP pairs with a range of thermodynamic and kinetic properties would provide a route to investigate the significance of the cooperative assembly (Fig. 2A).

**Figure 2.**
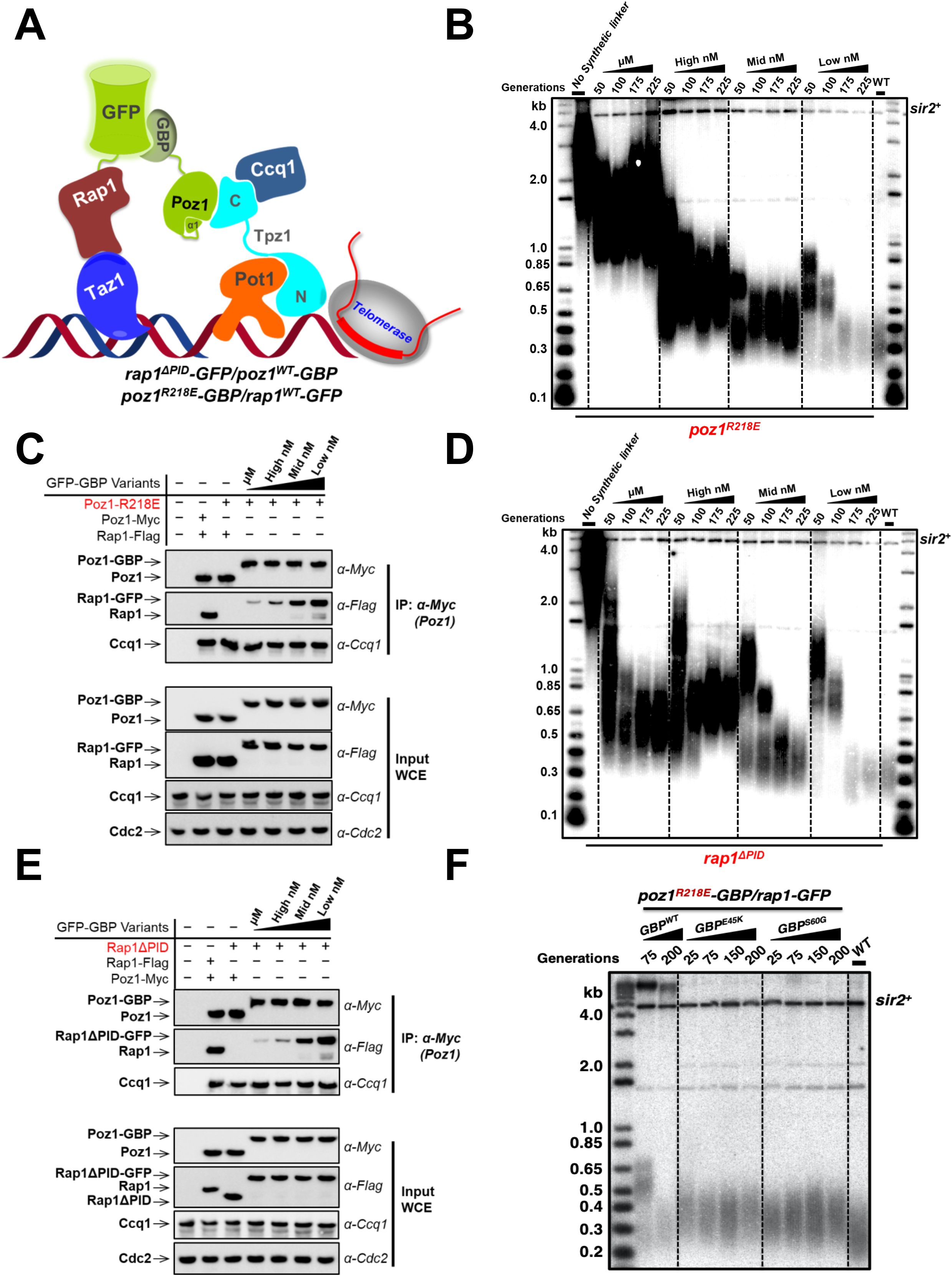
Slower disassociation rate rather than increased binding affinity is the key contribution of cooperative shelterin bridge assembly to telomere length regulation. (**A**) Schematic diagram of the synthetic *S. pombe* shelterin bridge with disrupted Rap1-Poz1 interaction. GFP is tagged to Rap1, and GBP variants are tagged to Poz1. (**B**) and (**D**) Telomere length analysis of indicated synthetic shelterin bridge strains in *poz1*^*R218E*^ (B) or *rap1*^*ΔPID*^ (D) background from successive re-streaks on agar plates via southern blotting. The telomere fragment is released from genomic DNA by ApaI digestion. Wild-type cells are denoted as WT. The mutants *poz1*^*R218E*^ (B) or *rap1*^*ΔPID*^ without synthetic shelterin bridge serve as controls and are denoted, “No Synthetic Linker”. *sir2*^*+*^ indicates an ApaI digested *sir2*^+^ gene fragment as the loading control. In this paper, the 1 kb plus marker from Life Technologies is used in all telomere length analysis. (**C**) and (**E**) Co-immunoprecipitation assays evaluating interactions among shelterin components with synthetic shelterin bridges in *poz1*^*R218E*^ (C) or *rap1*^*ΔPID*^ (E) background. Both Poz1-Rap1 and Poz1-Tpz1-Ccq1 interactions are measured. Cdc2 was shown as the loading control. Input: 1/30 of input WCE (whole-cell extract). (**F**) Telomere length analysis of *poz1*^*R218E*^ strains with GFP-*GBP*^*E45K*^ and GFP-*GBP*^*S60G*^ pairs as synthetic shelterin bridges, which have similar half-life as the native shelterin bridge.

We first assessed the consequence of using GFP-GBP variants to rescue telomere elongation in *poz1-R218E* background due to disruption of Poz1-Rap1 interaction within the shelterin bridge. In these strains, *poz1^+^* was tagged with GBP variants and Rap1 was tagged with GFP (Fig. 2A). As expected, *poz1-R218E* strain itself causes telomere massive elongation to ~ 3 kb due to loss of negative regulation from connected shelterin bridge. However, telomeres in *poz1-R218E* background with GFP-GBP variants linking *poz1-R218E* and *rap1^+^* showed various degrees of decreased length compared to the ~ 3 kb telomeres in *poz1-R218E* cells (Fig. 2B). Accordingly, the interactions among shelterin components were also rescued in a quantitative fashion as shown by co-immunoprecipitation experiments (Fig. 2C). As expected, the tighter the binding between GFP and GBP, the shorter the telomere length. Interestingly, Low nM GFP-GBP pair can rescue the telomere length to the wild-type level, whereas Mid nM GFP-GBP pair still have telomeres about 200 bp longer than the wild-type strain. The same outcome was also recapitulated in the GFP-GBP pair linked Rap1-Poz1 in the *rap1ΔPID* background (Fig. 2D and 2E), indicating that telomere length restoration is due to GFP-GBP pair-mediated linkage and is independent of the way that shelterin bridge is disrupted. It is worth noting that the native interface between Rap1 and Poz1-Tpz1 in the shelterin bridge has binding affinity close to Mid nM GFP-GBP pair (120 nM vs. 115 nM). However, our experiments indicate that similar affinity by itself cannot fully rescue the telomere length regulation defect. Instead, to fully rescue telomere length, synthetic shelterin bridge without cooperativity requires a GFP-GBP pair with ~110-fold higher binding affinity (K_d_^GFP-GBP^=1.06 nM vs. native shelterin bridge=120 nM) than that of the native shelterin bridge. This suggests that other advantages provided by cooperative shelterin bridge assembly, rather than thermodynamic affinity, plays a more important role in telomere length regulation.

We then compared the binding kinetics of synthetic shelterin bridges (GFP-GBP pairs) with that of Rap1 and Poz1-Tpz1 interaction. Based on its *k*_*off*_, and thus half-life (*t*_*1/2*_ = ln 2/*k*_*off*_), Poz1 and Rap1 interaction is intrinsically unstable with half of the complexes disassemble every 0.1 minute. Tpz1 stabilizes the Poz1-Rap1 interaction with an increased half-life of 6.42 min, over 60-fold increase. Among the four GFP-GBP pairs, only the Low nM GFP-GBP^WT^ pair offers longer half-life than the native shelterin bridge (70.4 min vs. 6.42 min). For Mid nM GFP-GBP^R36K^ pair, although its K_d_ is similar to that of the native shelterin bridge, its half-life (1.25 min) is about 5-fold shorter. The failure of Mid nM GFP-GBP^R36K^ pair to restore telomere length implies that binding kinetics (*k*_*off*_ or *t*_*1/2*_), rather than binding strength (K_d_), contributes more to telomere length regulation.

To further explore the contribution of shelterin bridge lifespan to telomere length regulation, we screened and obtained two more GFP-GBP variant pairs, which have more similar half-life *t*_*1/2*_ to the native shelterin bridge, GFP-GBP^E45K^ (*t*_*1/2*_ = 15.16 min) and GFP-GBP^S60G^ (*t*_*1/2*_ = 18.02 min), and 30-fold higher affinity (Fig. S1B). Indeed, when we introduced these two pairs of synthetic shelterin bridges to *poz1-R218E* strain background as before, we found that both of them can almost fully restore the telomere length (Fig. 2F), certainly to the comparable level as GFP-GBP^WT^. Thus, the kinetic lifespan of shelterin bridge assembly, rather than its thermodynamic affinity, plays a more important role in telomere length control. The extensively elongated lifespan of the shelterin bridge complex stemming from Tpz1-Poz1-Rap1 cooperative assembly might control the timespan of telomerase to telomere ends to regulate telomerase action. Therefore, cooperative assembly provides a “kinetic gateway” in shelterin bridge that controls the timespan of “open” and “closed” states of telomeres. Longer half-life of shelterin bridge (such as those in mid nM, high nM and μM synthetic shelterin bridges) leads to longer “open state” of the telomeres, thus providing more opportunities for telomerase to elongate telomeres.

### Rescuing of telomeric silencing only depends on the binding affinity within synthetic shelterin bridge

The complete linkage among shelterin components (Taz1-Rap1-Poz1-Tpz1-Ccq1) (Fig. 3A) is required to recruit CLRC to telomeric region for subtelomeric heterochromatin assembly, thus telomere silencing effect. To evaluate the function of cooperativity in shelterin-mediated heterochromatin assembly, we assessed silencing of *ura4^+^* reporter gene located adjacent to telomere region on a minichromosome. Consistent with previous studies, silencing of *TEL::ura4^+^* is defective in *rap1ΔPID* cells due to comprised shelterin bridge, resulting in cell lethality on +FOA plates. As expected, the defective silencing of *TEL::ura4^+^* is gradually alleviated in the stains with synthetic shelterin bridges. More cells are able to grow on +FOA plate along with stronger GFP-GBP variants interaction (Fig. 3B). Indeed, co-IP results also confirmed the restoration of the recruitment of CLRC by the shelterin complex, which was indicated by the rescued interaction between Rap1 and Clr4 with Mid nM GFP-GBP^R36K^ and Low nM GFP-GBP^wt^ synthetic shelterin bridges (Fig. 3C). Unexpectedly, although Low nM GFP-GBP^wt^ restored Rap1-Clr4 interaction much more than the Mid nM GFP-GBP^R36K^ did (over 20-fold, correlated with their K_d_), both of them can restore the telomeric silencing effect to a similar level (Fig. 3B). Therefore, different from telomere length regulation that requires slower dissociation kinetics enabled by the cooperative assembly, restoration of shelterin bridge to the wild-type binding affinity (Mid nM) is sufficient to rescue telomeric silencing. This result agrees with the passive recruitment role of shelterin in enriching CLRC methyltransferase complex onto the subtelomere regions for gene silencing, a process not dependent on time scale.

**Figure 3.**
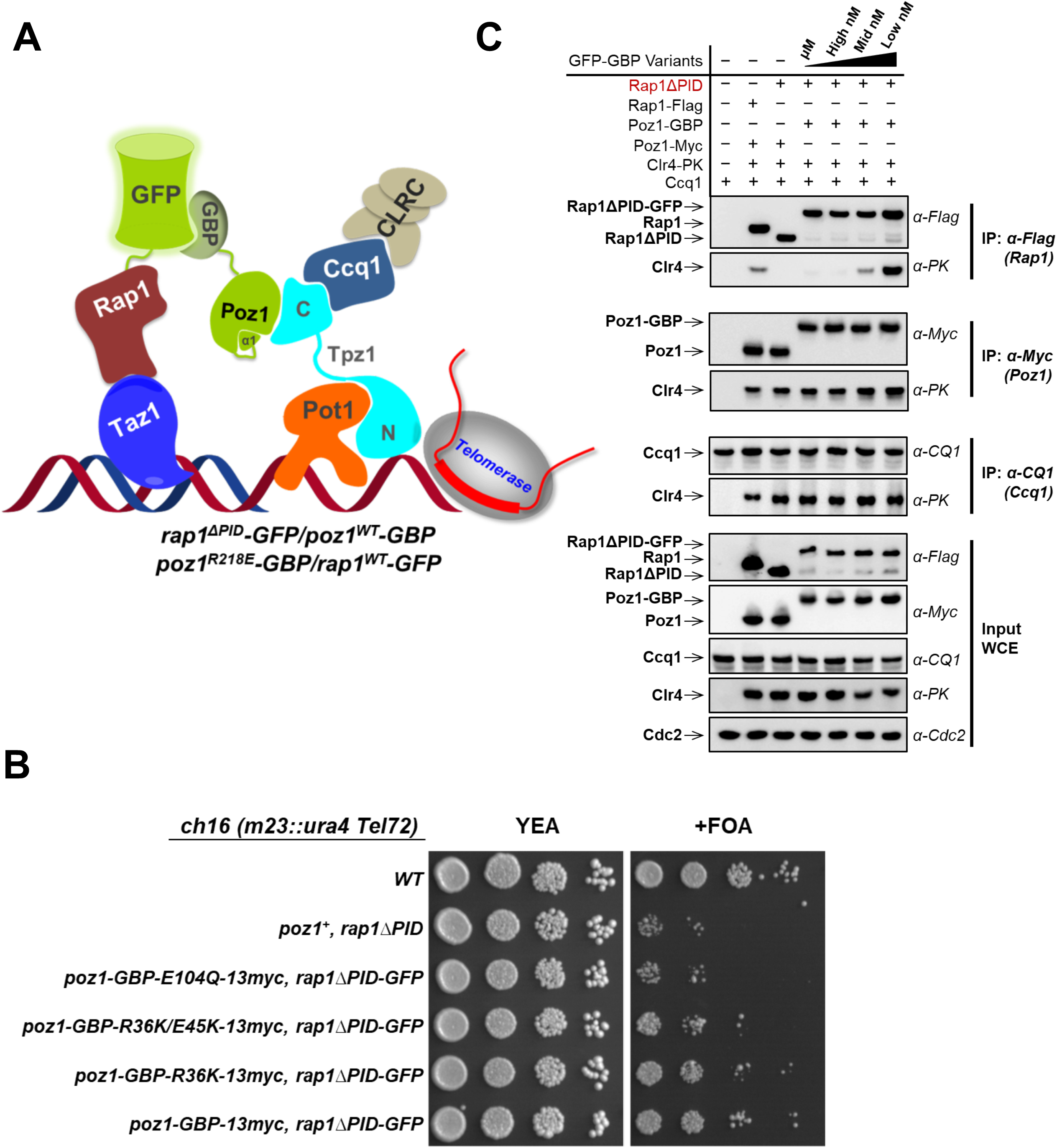
Shelterin-mediated telomeric silencing depends on the binding affinity within synthetic shelterin bridge. (**A**) Schematic diagram of the synthetic shelterin bridge recruiting CLRC complex to the telomere in *rap1^ΔPID^* background. (**B**) Tenfold serial dilution analyses of synthetic shelterin strains with *rap1^ΔPID^* grown on the indicated media to measure the expression of *TEL::ura4+*. (**C**) Co-immunoprecipitation assays evaluating synthetic shelterin bridge in CLRC complex recruitment for corresponding strains with *rap1^ΔPID^* background. Rap1-Clr4, Ccq1-Clr4, and Poz1-Clr4 interactions are measured.

### Conformational trigger in Poz1 is the key element for the “kinetic gateway”

In the process of cooperative shelterin bridge assembly, the very N-terminus of Poz1 (Poz1-NTD) have been shown to trigger the conformational changes in Poz1 upon Poz1-Tpz1 interaction (Kim et al., 2017). The conformational changes in Poz1 enhance Poz1-Rap1 interaction by increasing Poz1-Rap1 binding affinity and decreasing disassociation rate. Deletion of Poz1-NTD (Poz1ΔNTD) complete abolishes the high binding affinity binding between Poz1 and Rap1 even in the presence of Tpz1. As a result, *poz1ΔNTD* cells have drastically elongated telomeres. Interestingly, in *poz1ΔNTD* cells, Poz1-Rap1 and Poz1-Tpz1-Ccq1 interaction are both diminished, indicating essential role of the “conformational trigger” in regulating shelterin bridge and thus controlling telomere length. Therefore, we aim to assess whether the “conformational trigger” controls the “kinetic gateway”. Taking advantage of our GFP-GBP variant pairs, we fused the Low nM pair to Poz1-Rap1 and Poz1-Tpz1, respectively. Then, we assessed the shelterin bridge assembly in both settings via co-IP experiments. In the *poz1ΔNTD-GBP/rap1-GFP^wt^* background (Fig. 4A), we clearly observed Poz1-Rap1 interaction restored to the wild-type level (Fig. 4B). Intriguingly, Poz1-Tpz1-Ccq1 interaction was also partially restored (Fig. 4B). On the other hand, in the *poz1ΔNTD-GBP/tpz1-GFP^wt^* background (Fig. 4C), Poz1-Tpz1-Ccq1 interaction was fully rescued, but the synthetic linker had little effect on Poz1-Rap1 interaction (Fig. 4D). These results suggest that Poz1-NTD, the “conformational trigger”, play an essential role in shelterin bridge assembly by controlling Poz1-Rap1 interaction kinetics, which influences the Poz1-Tpz1-Ccq1 interaction of the bridge. Without the “conformational trigger”, even if Poz1 and Tpz1 are connected, Poz1 still cannot interact with Rap1. Agreeing with the co-IP results, whereas Low nM synthetic bridge in the *poz1ΔNTD-GBP/rap1-GFP^wt^* strain can almost rescue its telomere length to the wild-type level, synthetic bridge of the same strength in *poz1ΔNTD-GBP/tpz1-GFP^wt^* strain failed to restore the telomere length the to the same level (Fig. 4E). Not surprisingly, synthetic bridge of lower strength, such as Mid nM and μM GFP-GBP pairs, cannot restore the telomere length to the wild-type level either (Fig. S4).

**Figure 4.**
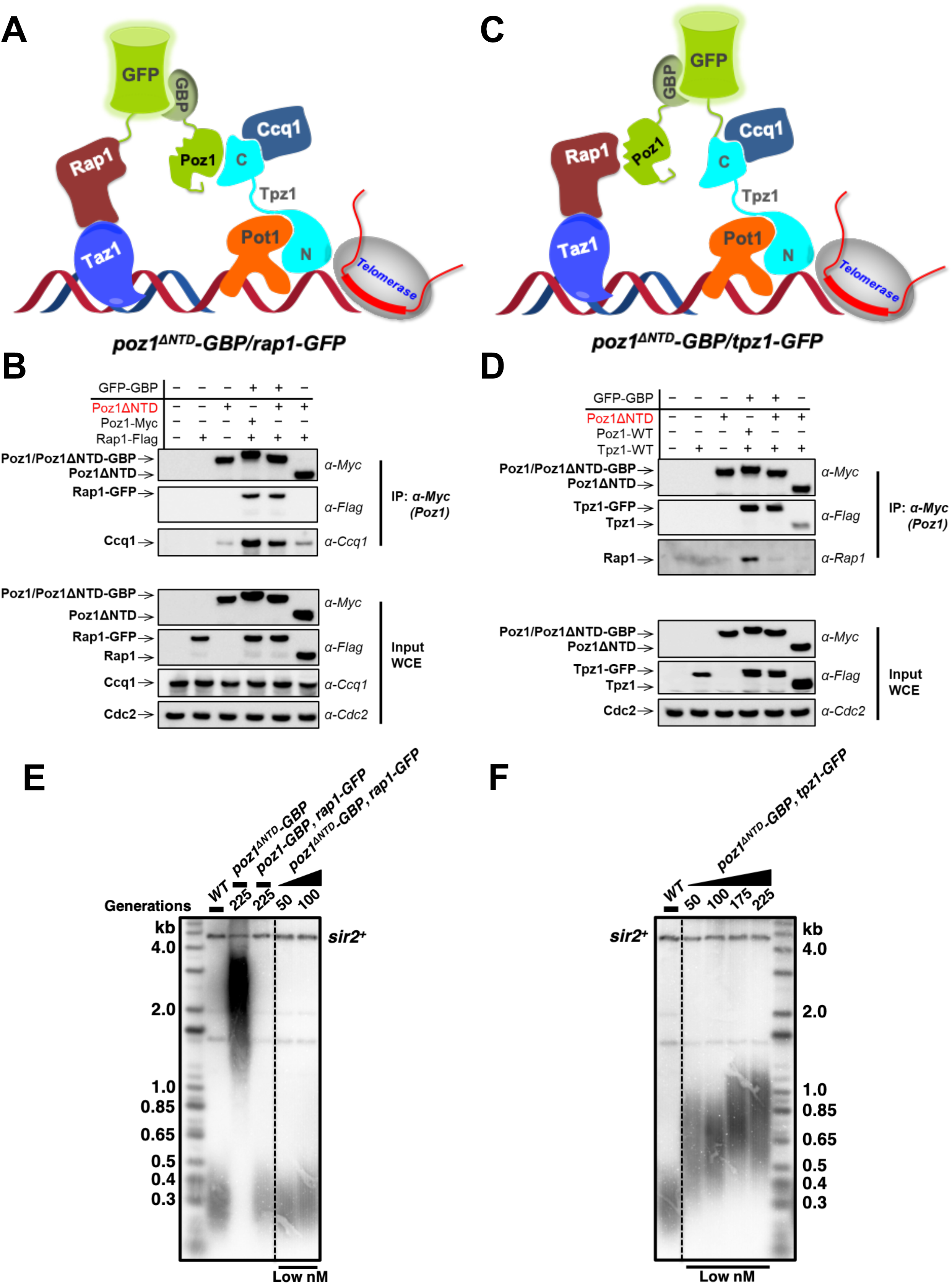
Conformational trigger in Poz1 is the key element for the “kinetic gateway”. (**A**) and (**C**) Schematic diagram of the synthetic *S. pombe* shelterin complex with conformational trigger in Poz1 deleted. Rap1 (A) or Tpz1 (C) is tagged with GFP, and Poz1 is tagged with GBP^WT^. (**B**) and (**D**) Co-immunoprecipitation assays evaluating the assembly of synthetic shelterin for the corresponding strains with *poz1^ΔNTD^*. Either Rap1-Poz1 (B) or Poz1-Tpz1 (D) is tethered via GFP-GBP pair. (**E**) and (**F**) Telomere length analysis of indicated synthetic shelterin strain in *poz1^ΔNTD^* background. Either Rap1-Poz1 (E) or Poz1-Tpz1 (F) is tethered via GFP-GBP pair.

## Discussion

As Richard Feynman said, “What I cannot create, I do not understand”. Inspired by this quote, we engineered *S. pombe* cells with synthetic shelterin bridges to carry out the telomere length regulation and telomeric silencing functions. This was enabled by the creation of GFP-GBP variant pairs with a wide range of thermodynamic and kinetic properties. Utilizing these synthetic shelterin bridges, we found that kinetic properties of the shelterin assembly, such as dissociation rate (half-life), has an unrecognized contribution to telomere length regulation. The intrinsic kinetic behavior of the shelterin assembly revealed in our study indicates its importance in collaborating with telomerase to elongate telomeres. Telomere lengthening is coupled to cell cycle-regulated events at telomere regions. In late S phase, when the DNA replication machinery completes most of the genome, Rad3^ATR^/Tel1^ATM^ are activated and phosphorylate the critical Thr93 residue in Ccq1 at telomeres, priming the telomere for telomerase recruitment. Then, telomerase holoenzyme, is recruited via two-pronged telomere-telomerase interfaces to the telomere by the cell cycle-regulated, phospho-Thr93-mediated Ccq1-Est1 and Trt1-Tpz1^TEL-Patch^ interactions (Chang et al., 2013). This intermediate telomerase recruitment complex further engages the telomerase core enzyme (Trt1-TER1) at the very 3’ end of the telomere for nucleotide additions. On the other hand, the substrate—telomeric DNA is also regulated. For elongation by telomerase, the very 3’ end of the telomere has to be in the extendible state. We found previously that the complete linkage between telomeric dsDNA binder and ssDNA binder controls telomeres in the non-extendible state. Permanent breakage of the linkage leads to drastically elongated telomeres. In this study, we demonstrated the correlation between the life span of shelterin bridge and telomere length. If the life span of the bridge is too short, for example with *t*_*1/2*_ less than 2 min (in the cases of Mid nM to μM synthetic bridges, and *poz1*^*ΔNTD*^), the bridge would be mostly in the open state during the late S phase (20-40 min), thus providing telomerase high percentage of extendible telomeres to elongate. This loss of “open” and “close” state control on the telomere (substrate) side leads to massive elongation of telomeres. In contrast, for native shelterin bridge assembled with cooperativity, the life span of the bridge t_1/2_ is 6.42 min (Fig. 5), which provides optimal percentage of open telomeres for telomerase to elongate during the late S phase. On top of cell cycle-regulated telomerase recruitment, our study adds an additional layer of temporal regulation of telomere elongation through the kinetic gateway intrinsic to shelterin complex assembly.

**Figure 5.**
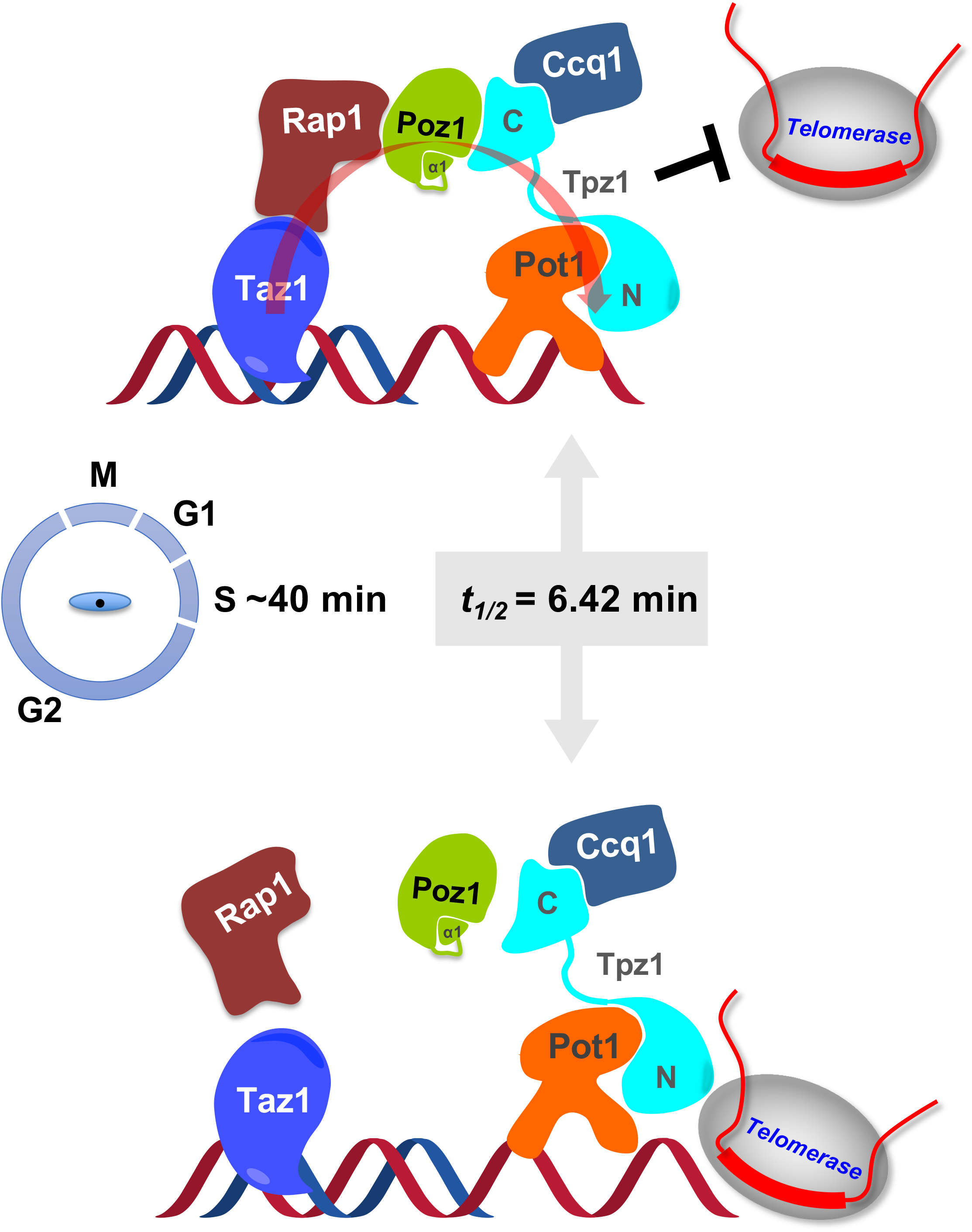
Model of how kinetics of shelterin bridge assembly and disassembly controls telomere elongation. Spatiotemporal regulation of telomerase-mediated telomere elongation is coupled to telomere states controlled by shelterin complex. Cooperative assembly confers kinetic properties on the shelterin bridge allowing disassembly to function as a molecular timer, regulating the switching of the telomere open and closed states to achieve a defined species-specific length range.

Interestingly, for the role of shelterin bridge in establishing heterochromatin and telomeric silencing, Mid nM and Low nM synthetic bridge shows almost no difference in restoring the telomeric silencing effect. This is different from telomere length regulation that requires slower dissociation kinetics enabled by the cooperative assembly. This is most likely due to the passive recruitment role of shelterin in enriching CLRC methyltransferase complex onto the subtelomere regions for gene silencing, a process not dependent on time scale, but determined by the critical concentration of CLRC enriched by the shelterin bridge to the telomeric and subtelomeric regions. Clearly, distinctive kinetic and thermodynamic properties of shelterin bridge contribute accordingly to the biological processes it participates in.

## Materials and Methods

### Yeast Strains, Gene Tagging, and Mutagenesis

Fission yeast strains used in this study are listed in supplemental Table S1. Wild type tagging strains and single mutant strains were constructed by one-step gene replacement of the entire ORF with the C-terminus epitope-tags followed by selectable markers. The pFA6a plasmid modules were used as templates for epitope-tags and selectable markers. The GFP-3Flag and GBP-13myc tagging plasmids were engineered from pFA6a-GFP-KanMX6 and pFA6a-GBP-hphMX6 plasmids through mutagenesis PCR, respectively. Double mutant strains were produced by mating, sporulation, dissection, and selection followed by PCR verification of genotypes. All mutations were confirmed by DNA sequencing (Eton, San Diego, CA). For serial dilution plating assays, 10-fold dilutions of a log-phase culture were plated on the indicated medium and grown for 4 d at 30°C.

### Protein expression and purification

Target proteins were subcloned into modified pET28a vector containing either 10His-Smt3 tag or the Avi-6His-SUMO tag (Zhao et al., 2016). Plasmids were transformed into Rosetta-BL21 (DE3) cells for protein expression, which was induced with 0.4 mM IPTG for 4 h at 30 °C. For biotinylated proteins, a pBirAcm vector was co-transformed. Cells were disrupted by sonication in lysis buffer (25mM Tris-HCl at pH 8.0, 350 mM NaCl, 15 mM imidazole, 5 mM b-mercaptoethanol, 1 mM PMSF, 2 mM benzamidine). The supernatant was cleared with centrifuge and incubated with Ni-NTA (Qiagen) resin for 1 h. The elution buffer (lysis buffer plus 300 mM imidazole) was used to elute the proteins from resin. Then the proteins were further purified with gel filtration columns.

### Biolayer Interferometry (BLI) and Kinetics Measurement

The BLI experiment and data analysis were performed as previously described. Briefly, biotinylated GBP proteins, including WT and mutants, were loaded on Streptavidin biosensor tips, followed by quenching free streptavidin with biocytin. GFP was then added to measure association and dissociation rate. Data was analyzed with ForteBio Data analysis version 9.0 and fitted with global/1:1 binding model. Equilibrium dissociation constant (K_d_) and association (*k*_*on*_) and dissociation (*k*_*off*_) rate constants were calculated directly from software; half-life (*t*_*1/2*_) was then further calculated *k*_*off*_.

### Telomere Length Analysis

The telomere length of each strain was analyzed as previously described (Liu et al., 2015). Briefly, cells were successively cultured on YEAU plates and genomic DNA from each generation was prepared from 5 ml liquid culture inoculated from plates. The telomeric fragments were released by ApaI (NEB) digestion and separated on 1.8% agarose gels. Southern blots with both telomeric and *sir2^+^* probes were visualized using Typhoon scanner.

### Co-Immunoprecipitation

The indicated strains were cultured in 50 ml YEAU and harvested at log phase. Cell pellets were then washed and cryogenically disrupted with FastPrep MP with two pulses (60 sec) of bead-beating in ice-cold lysis buffer (50 mM HEPES at pH 7.5, 140 mM NaCl, 15 mM EGTA, 15 mM MgCl_2_, 0.1% NP40, 0.5 mM Na_3_VO_4_, 1 mM NaF, 2 mM PMSF, 2 mM benzamidine, Complete proteinase inhibitor [Roche]). After clearing with centrifuge, the protein concentration was measured via Bradford assay and adjusted to 12 mg/ml. Anti-Flag M2 affinity gel (Sigma), anti-Myc (9E10 from Santa Cruz Biotechnology) or anti-Ccq1 rabbit serum plus IgG beads (Roche) was used for immunoprecipitation, followed by eluting with 30 μl 0.1 M glycine (pH2.0) at room temperature for 10 min. The elute was immediately neutralized with 2μl 2 M Tris-HCl, pH 8.0. SDS-PAGE (8%) and western blotting using monoclonal anti-Flag (M2-F1804, from Sigma), monoclonal anti-Myc (from Covance), monoclonal anti-PK (ab27671 from Abcam), anti-Ccq1 rabbit serum, or anti-Cdc2 (y100.4, from Abcam) were performed to detect protein-protein interaction as indicated.

## Acknowledgements

We thank Fuyuki Ishikawa, Toru Nakamura, Julie Cooper, Virginia Zakian, Pingwei Li, and Takashi Toda for providing plasmids and strains, Craig Kaplan and Peter Kaiser for comments on the manuscript and helpful discussions. This work was supported by an NIH grant R01GM098943 and an American Cancer Society Research Scholar Grant RSG-16-041-01-DMC to F. Q. as well as R35GM126910 to S. J.

## Author Contributions

F. Q. conceived, designed, and supervised the study; J. L. made various constructs, purified the proteins, and performed biochemical experiments. J. L. and J. K. analyzed the biochemical data, and J. K. made protein structure figures. J. L. and X. H. performed genetic experiments and analyzed the data. K. B. and S. J. performed telomeric silencing assays. J. L., J. K., K. B. and X. H. prepared the figures. F. Q. and J. L. wrote the manuscript with input from S. J., J. K. and X. H.

---

**Figure S1. (Related to Figure 1).**
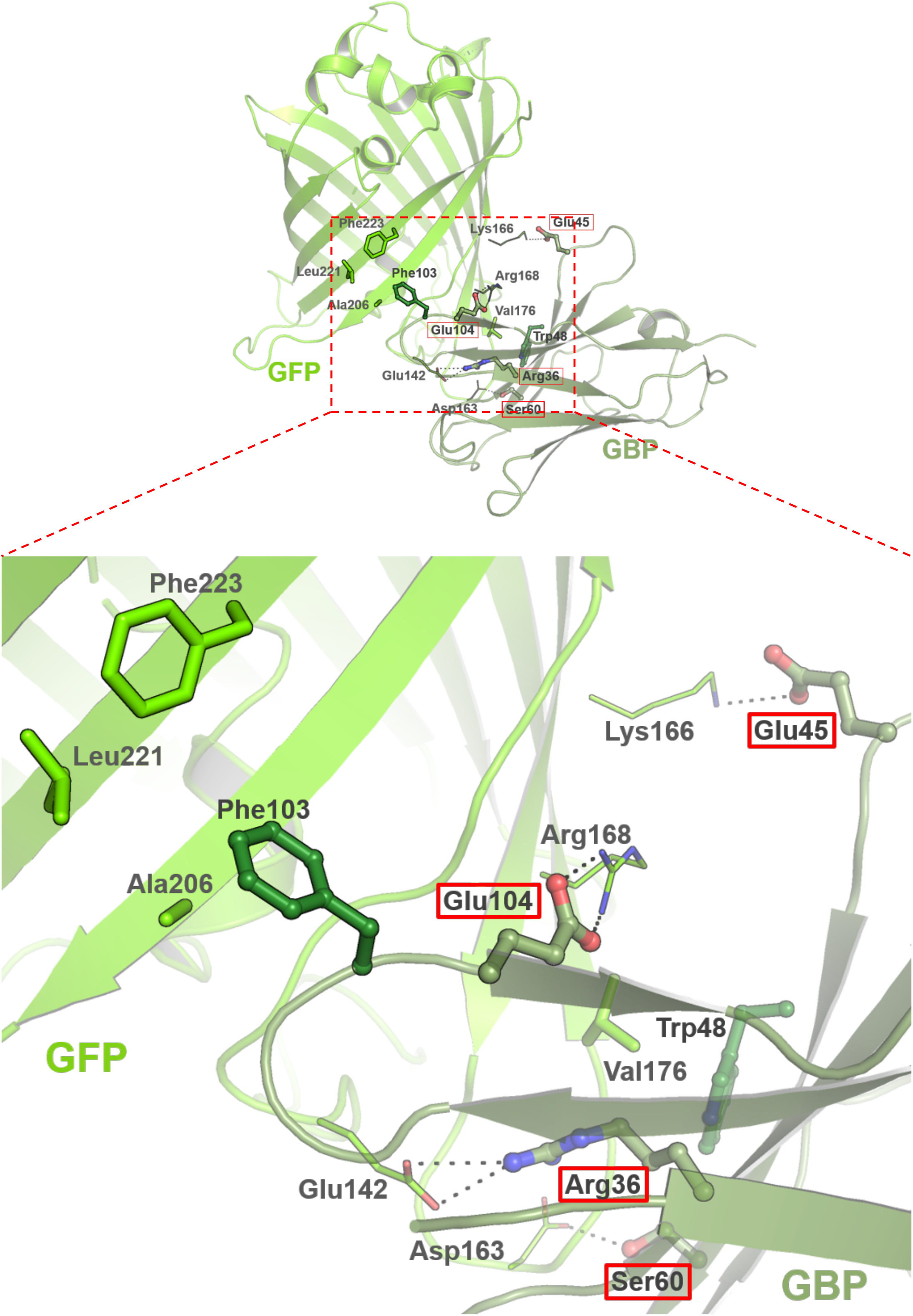
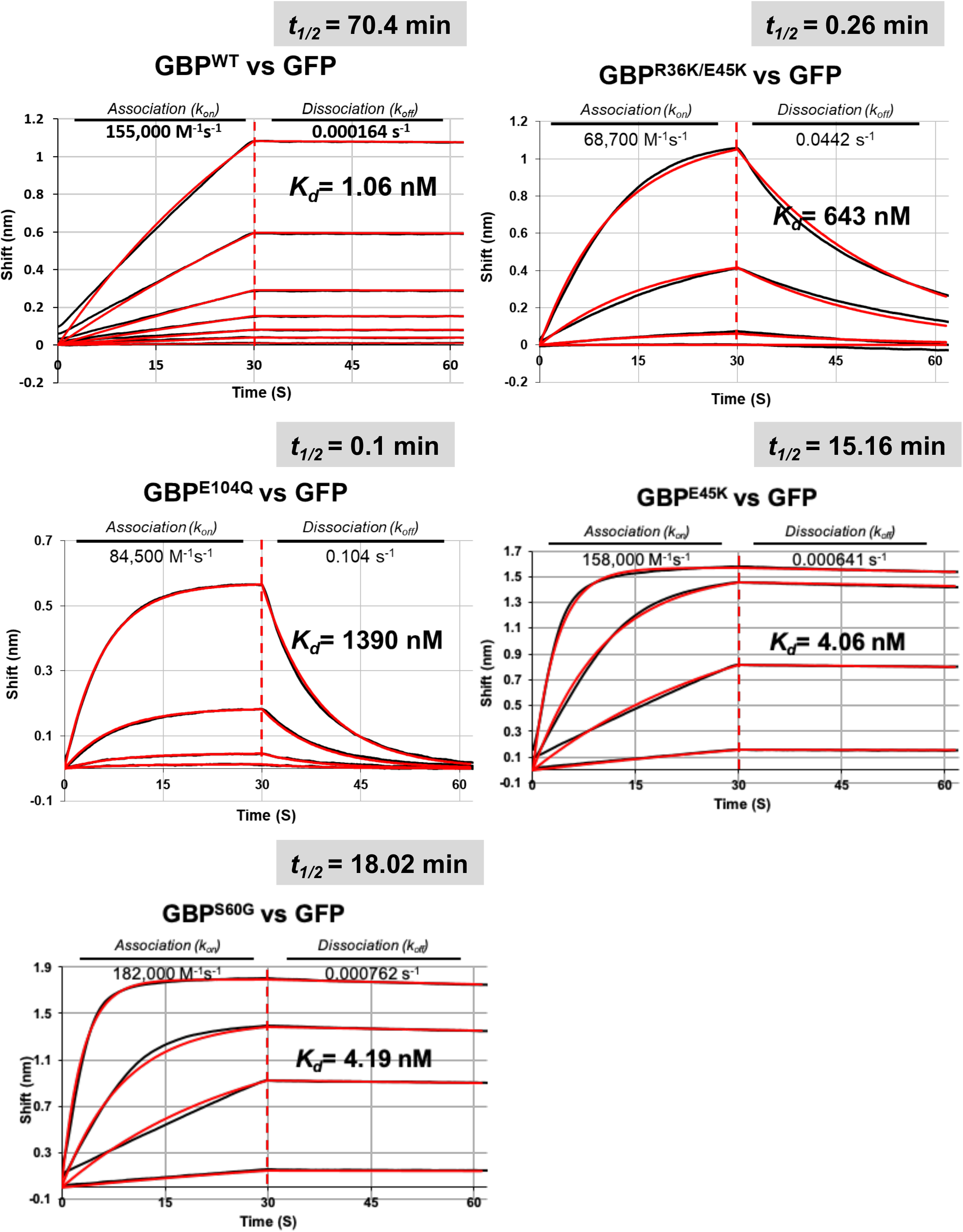
(**A**) Crystal structure of GFP-GBP complex. The interface is zoomed in for close-up view. (**B**) Bio-layer interferometry (BLI) sensorgrams measuring dissociation and association events in real time between GFP and GBP variants using Octet red96.

**Figure S2. (Related to Figure 2).**
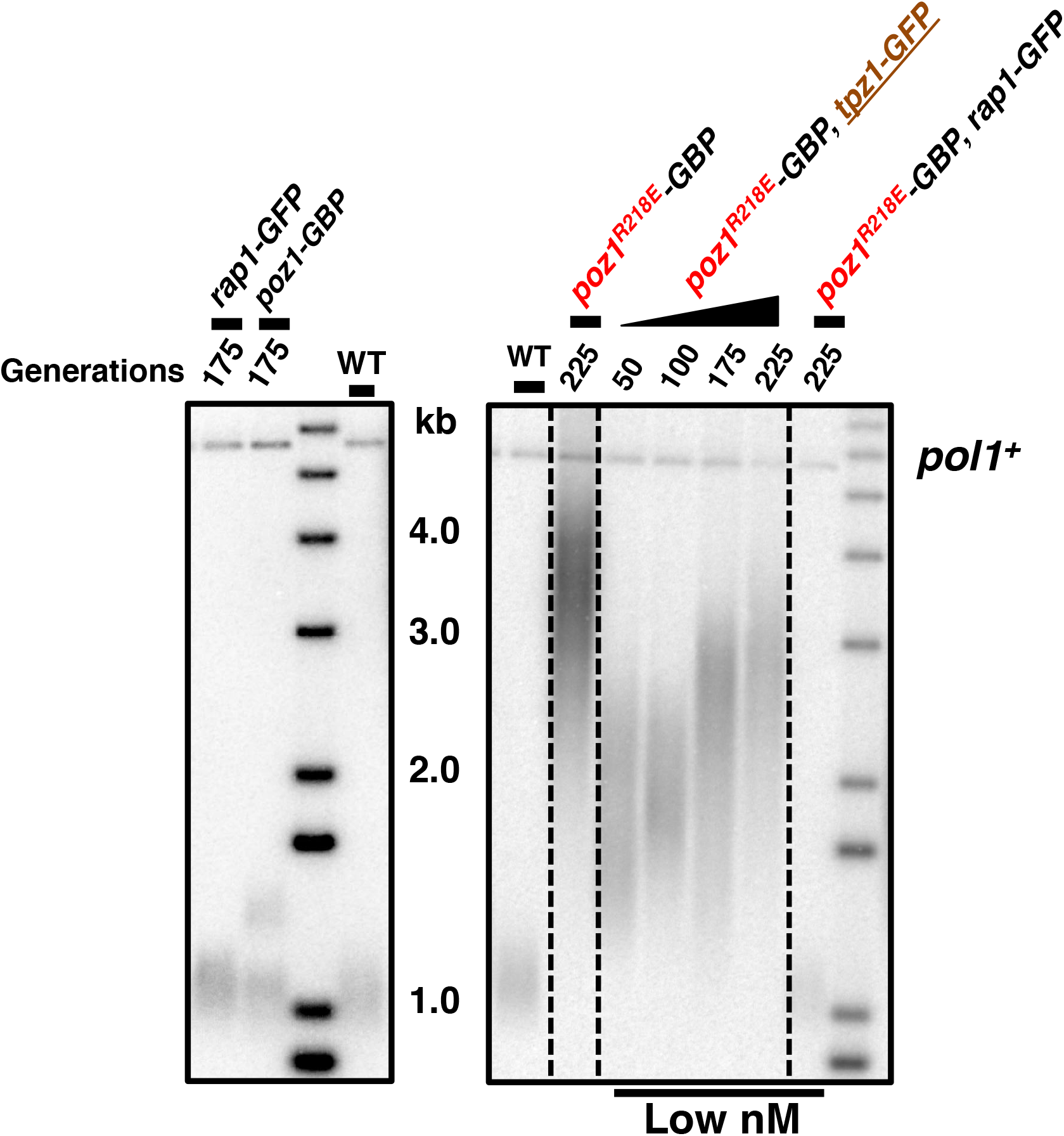
Tagging *rap1*^+^ and *poz1*^+^ with GFP and GBP, respectively, has no effect on telomere length maintenance (*left*). Pairing *tpz1*^+^ and *poz1*^+^ via GFP-GBP in *poz1*^*R218E*^ background cannot resort telomere length to WT level (*right*). Telomere fragments were released from EcoRI digested genomic DNA with *polI*^+^fragments serving as an internal loading control.

**Figure S4. (Related to Figure 4).**
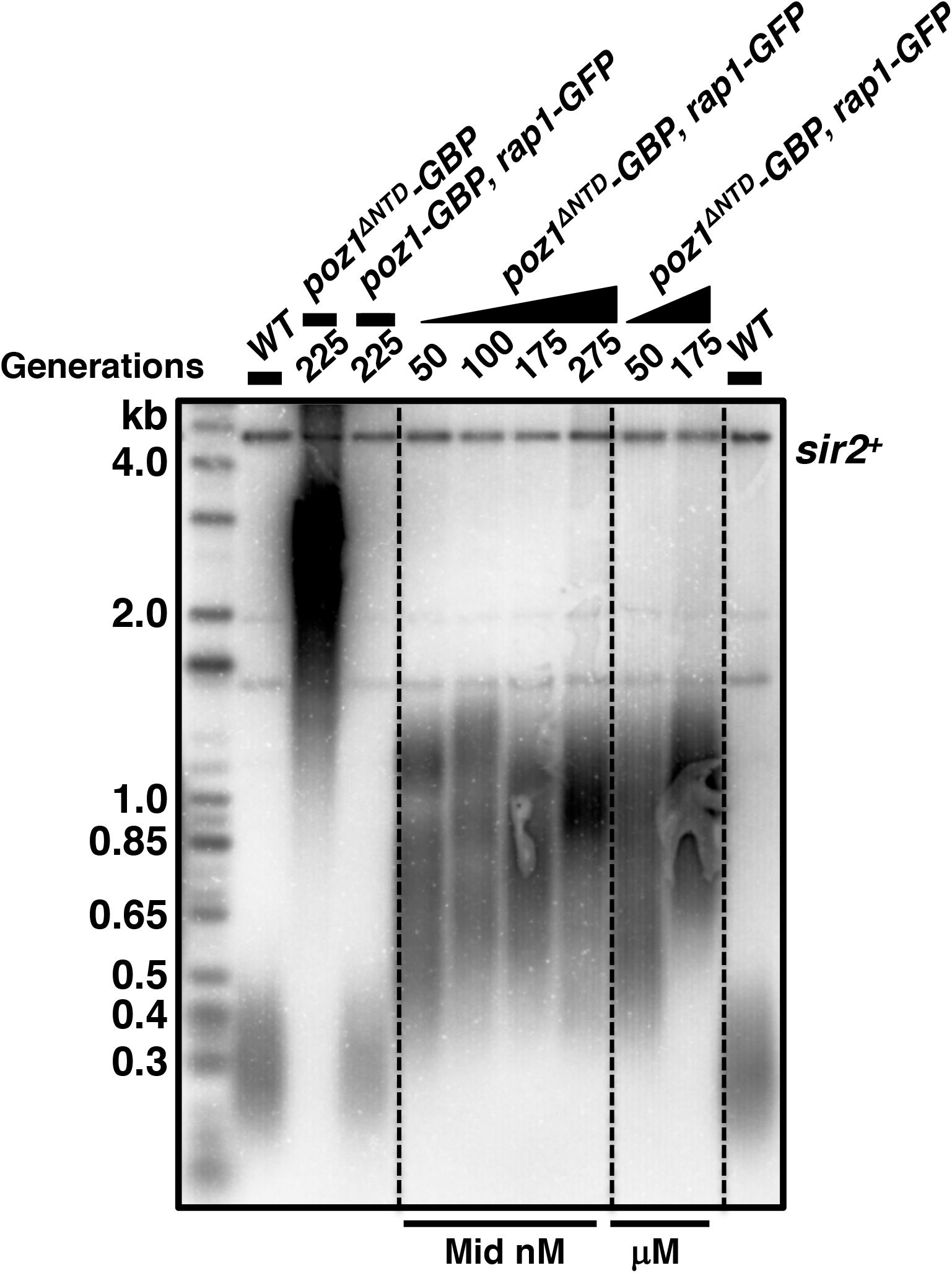
Mid nM or μM GFP-GBP linker fails to rescue telomere phenotype in loss of conformational trigger background (*poz1*^*ΔNTD*^).

**Supplemental Table S1.**
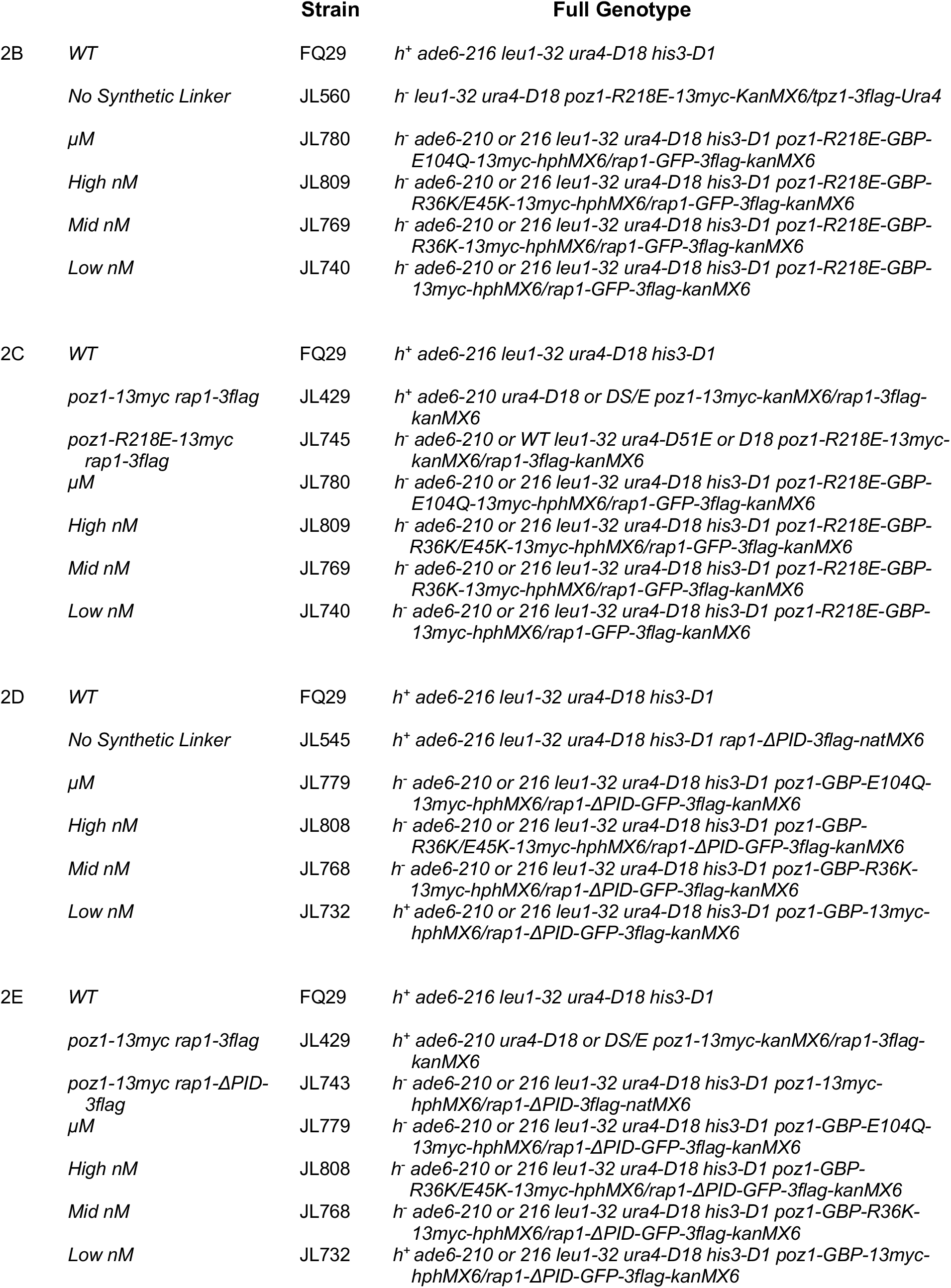
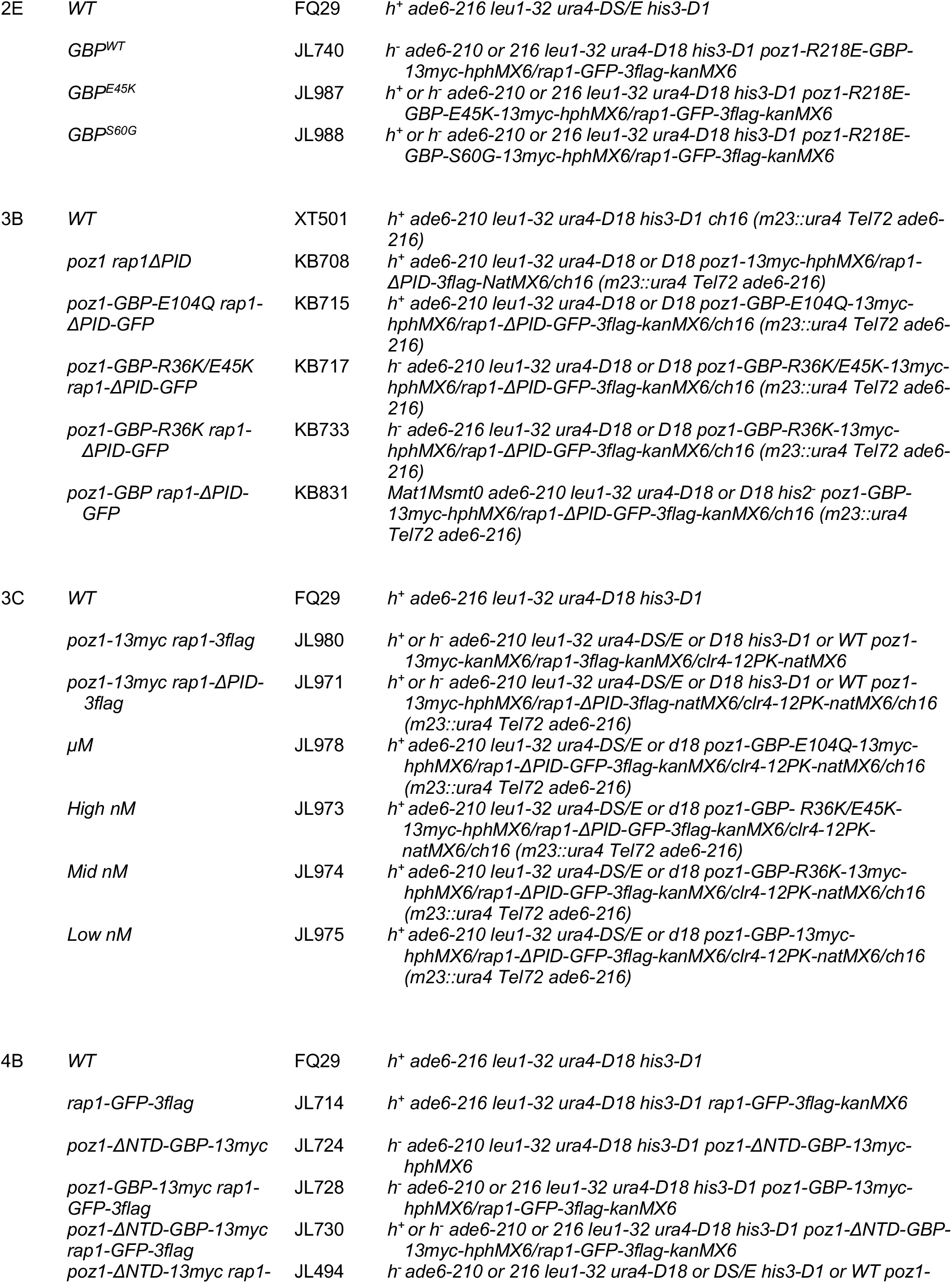
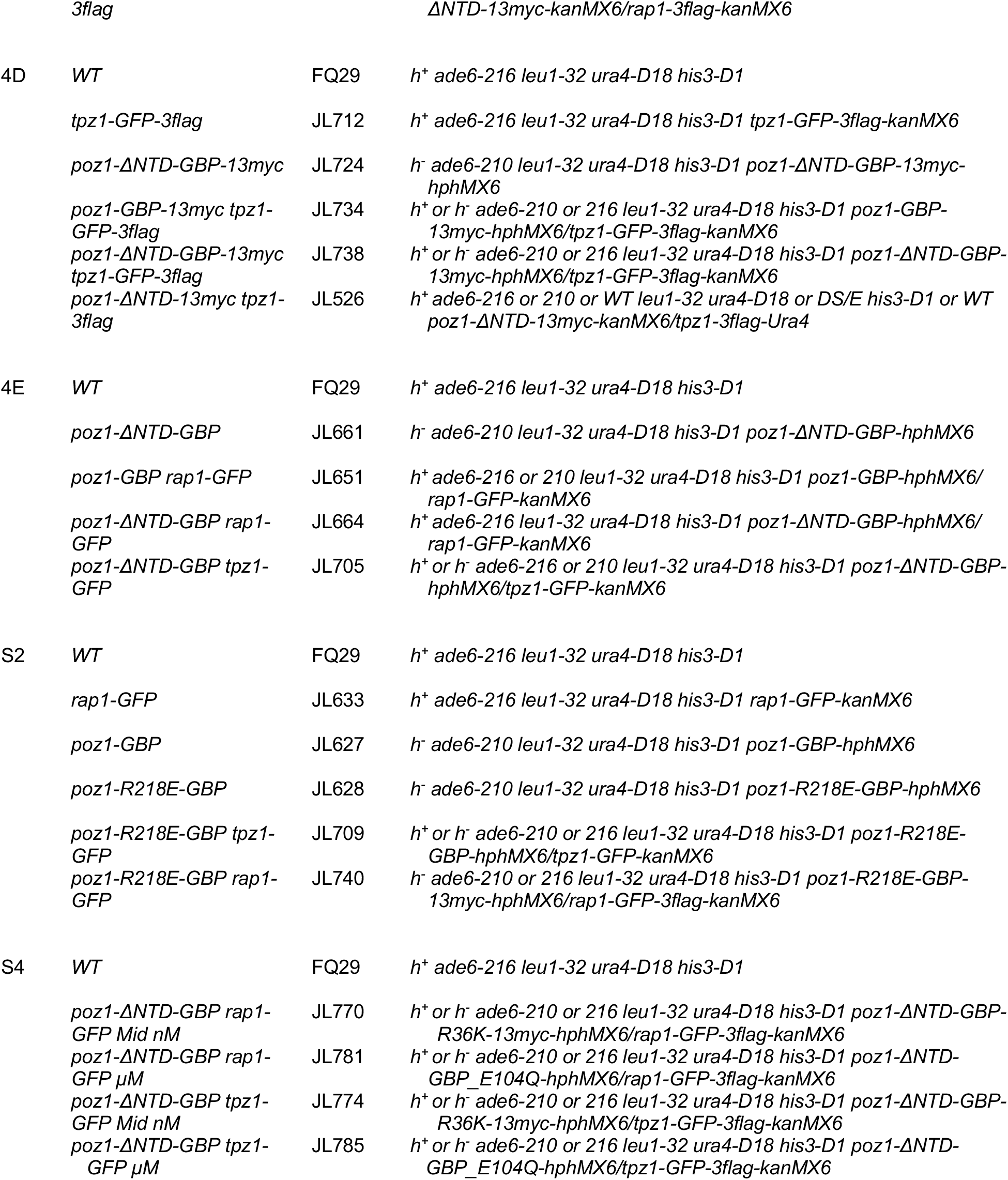
Fission yeast strains used in this study.

**Table S2.** Plasmids used in this study.

| Plasmid | Description | Restriction sites used | Primers | Source |
| --- | --- | --- | --- | --- |
| pRARE | *E. coli* Rosetta™ (DE3) plasmid, encodes rare tRNAs, Cam<sup>R</sup> |  |  | EMD Millipore |
| pET28a-His-smt3-GFP | Target sequence with N-terminal (His-tag)-smt3-Ulp1 cleavage site in pET28a vector | BamH I/Xho I | See Table SX |  |
| pET28a-Avi-His-smt3-GBP | Target sequence with N-terminal (Avi-tag)-(His-tag)-smt3-Ulp1 cleavage site in pET28a vector | BamH I/Xho I | See Table SX |  |
| pET28a-Avi-His-smt3-GBP-E104Q | Target sequence with N-terminal (Avi-tag)-(His-tag)-smt3-Ulp1 cleavage site in pET28a vector | BamH I/Xho I | See Table SX |  |
| pET28a-Avi-His-smt3-GBP-R36K/E45K | Target sequence with N-terminal (Avi-tag)-(His-tag)-smt3-Ulp1 cleavage site in pET28a vector | BamH I/Xho I | See Table SX |  |
| pET28a-Avi-His-smt3-GBP-R36K | Target sequence with N-terminal (Avi-tag)-(His-tag)-smt3-Ulp1 cleavage site in pET28a vector | BamH I/Xho I | See Table SX |  |
| pET28a-Avi-His-smt3-GBP-E45K | Target sequence with N-terminal (Avi-tag)-(His-tag)-smt3-Ulp1 cleavage site in pET28a vector | BamH I/Xho I | See Table SX |  |
| pET28a-Avi-His-smt3-GBP-S60G | Target sequence with N-terminal (Avi-tag)-(His-tag)-smt3-Ulp1 cleavage site in pET28a vector | BamH I/Xho I | See Table SX |  |
| pBirAcm | pACYC184 background IPTG inducible biotin Ligase vector, Cam<sup>R</sup> |  |  | avidity |
| pFA6a-GFP-3 Flag-kanMX6 | Endogenous tagging with C-terminal GFP-3Flag in pFA6a vector |  | See Table SX |  |
| pFA6a-GBP-13 myc-hphMX6 | Endogenous tagging with C-terminal GBP-13myc in pFA6a vector |  | See Table SX |  |
| pFA6a-GBP-E104Q-13myc-hphMX6 | Endogenous tagging with C-terminal GBP-E104Q-13myc in pFA6a vector |  | See Table SX |  |
| pFA6a-GBP-R36K/E45K-13 myc-hphMX6 | Endogenous tagging with C-terminal GBP-R36K/E45K-13myc in pFA6a vector |  | See Table SX |  |
| pFA6a-GBP-R36K-13myc-hp hMX6 | Endogenous tagging with C-terminal GBP-R36K-13myc in pFA6a vector |  | See Table SX |  |
| pFA6a-GBP-E45K-13myc-hp hMX6 | Endogenous tagging with C-terminal GBP-E45K-13myc in pFA6a vector |  | See Table SX |  |
| pFA6a-GBP-S60G-13myc-hp hMX6 | Endogenous tagging with C-terminal GBP-S60G-13myc in pFA6a vector |  | See Table SX |  |

